# Effect of FMRFamide on voltage-dependent currents in identified centrifugal neurons of the optic lobe of the cuttlefish, *Sepia officinalis*

**DOI:** 10.1101/2020.09.29.318691

**Authors:** Abdesslam Chrachri

**Affiliations:** University of Plymouth, Dept of Biological Sciences, Drake Circus, Plymouth, PL4 8AA, UK and the Marine Biological Association of the UK, Citadel Hill, Plymouth PL1 2PB, UK

**Keywords:** cephalopod, voltage-clamp, potassium current, calcium currents, sodium current, FMRFamide

## Abstract

Whole-cell patch-clamp recordings from identified centrifugal neurons of the optic lobe in a slice preparation allowed the characterization of five voltage-dependent currents; two outward and three inward currents. The outward currents were; the 4-aminopyridine-sensitive transient potassium or A-current (I_A_), the TEA-sensitive sustained current or delayed rectifier (I_K_). The inward currents were; the tetrodotoxin-sensitive transient current or sodium current (I_Na_). The second is the cobalt- and cadmium-sensitive sustained current which is enhanced by barium and blocked by the dihydropyridine antagonist, nifedipine suggesting that it could be the L-type calcium current (I_CaL_). Finally, another transient inward current, also carried by calcium, but unlike the L-type, this current is activated at more negative potentials and resembles the low-voltage-activated or T-type calcium current (I_CaT_) of other preparations.

Application of the neuropeptide FMRFamide caused a significant attenuation to the peak amplitude of both sodium and sustained calcium currents without any apparent effect on the transient calcium current. Furthermore, FMRFamide also caused a reduction of both outward currents in these centrifugal neurons. The fact that FMRFamide reduced the magnitude of four of five characterized currents could suggest that this neuropeptide may act as a strong inhibitory agent on these neurons.

**Summary:** FMRFamide modulate the ionic currents in identified centrifugal neurons in the optic lobe of cuttlefish: thus, FMRFamide could play a key role in visual processing of these animals.

## Introduction

Among invertebrates, cephalopods are considered to have an extremely well-developed eye and a centralized brain (Williamson and Chrachri, 2004), their retina lacks the vertebrates equivalent of bipolar, amacrine, ganglion cells, etc. and therefore there is very little visual processing within their retina which instead take place in the optic lobe. Their retina contains only photoreceptors and supporting cells and it has been demonstrated that there are some interconnection between the photoreceptors (Yamamoto et al., 1965; Yamamoto and Takasu, 1984). There is also an extensive efferent innervation of the retina coming from the inner granular layer of the cortex of the optic lobe (Young, 1971 and 1974). These efferents are the axons of the centrifugal neurons that have been demonstrated to be involved in the regulation of the size of the receptive fields (Tasaki et al., 1982) and the control of the screening pigment migration (Gleadall et al., 1993). This is still relatively simple compare to the vertebrate retina.

Octopus seem to have a kind of camera eye with an iris and adjustable lens similar to those of vertebrates, the retina consists of a single layer of photoreceptor cells, and the optic lobe constitutes the center for visual analysis (Young, 1962). It has also been suggested that the centrifugal neurons in the optic lobe project towards the photoreceptors in the retina (Lund, 1979; Saidel, 1979). Although there is little data about the neuromodulators contained in the centrifugal neurons, the presence of several neurotransmitters and possible neuromodulators have already been observed in the optic lobe (Cornwell et al., 1993; Di Cosmo and Di Cristo, 1998; Kito-Yamashita et al., 1990; Sasayama et al., 1991; Suzuki and Yamamaoto, 2000 and 2002).

The neuropeptide FMRFamide (Phe-Met-Arg-Phe-NH_2_) and similar molecules which are collectively referred to as FMRFa-related peptides (FaRPs) first discovered in molluscs (Price and Greenberg, 1977 and 1989) are conserved throughout the animal phyla (Walker et al., 2009). They are abundant in both vertebrate and invertebrate nervous systems (Espinoza et al., 2000; Dockray et al., 1983; O’Donohue et al., 1984; Sorenson et al., 1984; Schneider and Taghert, 1988; Greenberg and Price, 1992; Nelson et al., 1998). In these organisms, FMRFa-like neuropeptide act as neurotransmitters and neuromodulators. In mammals, FMRFamide induces a variety of physiological effects, including alterations in blood pressure, respiratory rate, glucose-stimulated insulin release, and behavior (Mues et al., 1982; Sorenson et al., 1984; Raffa et al., 1986; Thiemermann et al., 1991; Muthal et al., 1997; Nishimura et al., 2000; Askwith et al., 2000). In cephalopods, it has been demonstrated that FMRFa is involved in the control of egg laying in *Sepia officinalis* (Henry et al., 1999) as well as the modulation of L-type calcium currents in heart muscle cells of squid (Chrachri et al., 2000) and that of both the excitatory and inhibitory postsynaptic currents in optic lobe neurones of cuttlefish (Chrachri and Williamson, 2003). In 1997, Loi and Tublitz reported the isolation and characterization of a full-length FaRP cDNA from the brain of cuttlefish, *Sepia officinalis.* The presence of FMRFa-like immunoreactivity in the optic lobe of both octopus (Suzuki et al., 2002) and cuttlefish (Chrachri and Williamson, 2003) has been reported indicating a putative neurotransmitter or neuromodulator role for this neuropeptide. Furthermore, receptor binding studies with squid optic lobes have identified G-protein associated FMRFa binding sites (Chin et al., 1994). Although there have been reports of the ability of this neuropeptide to modulate the activity of some cephalopod muscles (Loi and Tublitz, 2000; Chrachri et al., 2000) and to potentiate the activity at the squid giant synapse (Cottrell et al., 1992). However, there is less understanding of its central function (Chrachri and Williamson, 2003). In metazoans, different FMRFamide-Like Peptides (FLP) types may exist in the same species, and the same FLP type may occur in various species (Walker et al. 2009). Recently, a full-length cDNA sequence of an FMRFamide gene isolated from the cuttlefish, *Sepia pharaonis* was cloned which shares 93% and 92% similarity with two cuttlefish species, *Sepiella japonica and Sepia officinalis* (Li et al., 2018; Zhu et al., 2020).

The mechanisms by which these FaRPs act are not yet fully understood. However, it has been demonstrated that they either act directly on the membrane conductances of excitable cells (Cottrell et al., 1984; Colombaioni et al., 1985; Belkin and Abrams, 1993) or through second messenger system (Brezina, 1988; Chiba et al., 1992; Raffa and Stone, 1996; Chrachri et al., 2000). Furthermore, it has been also shown that FMRFa can also activate directly one or more types of ligand-gated channels (Cottrell et al., 1990; Chin et al., 1994; Green et al., 1994).

Although the morphology of the centrifugal neurons of the optic lobe of cephalopod have been studied using both Golgi and cobalt chloride staining (Young, 1974; Saidel, 1979), there is no information on the types of ionic currents present in cephalopod centrifugal neurons. This report provides the first comprehensive study of the voltage-activated whole-cell membrane currents present in identified centrifugal neurons; we have characterized five separate ionic currents in these neurons; two outward currents and three inward currents and studied their modulation by the neuropeptide FMRFamide.

## Material and methods

### Preparation of slices

Slices were prepared from both male and female cuttlefish, *Sepia officinalis.* Animals were anaesthetized with 2% ethanol and sacrificed by decapitation. The optic lobe was rapidly removed and placed in ice-cold Ca^2+^-free artificial sea water (ASW). Transverse slices of around 300 μm were cut with a vibratome (Campden Instruments, Loughborough, UK). Slices were kept in a storage chamber in the fridge for up to 1 h before recording.

### Electrophysiological recordings

Individual slices were transferred to the recording chamber, they were fully submerged and superfused with oxygenated ASW at a rate of 2-3 ml/min. Centrifugal neurons were visually identified using an Olympus BX50WI upright microscope with a x40 water immersion objective lens and equipped with infrared illumination and video enhanced visualization system consisting of a CCD camera (C7500) and its controller (C2741-90, Hamamatsu Photonics Ltd., Hertfordshire, UK).

Identified centrifugal neurons were studied at room temperature (18-20 °C) with the whole-cell patch recording techniques (Hamill et al., 1981). Recordings were made using an Axopatch amplifier (200A, Molecular devices, San Jose, CA 95134, USA) controlled by PClamp8 software (Molecular devices) for data acquisition, analysis and storage. Ionic currents were sampled at a rate of 10 kHz and were low-pass filtered at 2 kHz. The pipette series resistance was electronically compensated, as far as possible, to give voltage errors of only few mV at peak current levels. The capacitance current response to a −10 mV voltage step, from a holding potential of −60mV, was recorded for each neuron and the access resistance calculated. Liquid junction potential was calculated using the liquid junction calculator provided by the PClamp software. Patch electrodes with tip resistance of 4.0–6.0 MΩ were pulled using a horizontal puller (Sutter Model P-97, Novato, CA, USA) from soda glass capillaries (Intracel, 1.5 mm o.d., 0.86 mm i.d.). These electrodes were filled with either Aspartate or CsCl_2_ based internal solution.

### Solution and drugs

The artificial sea water contained (in mM): 430 NaCl; 10 KCl; 10 CaCl_2_; 30 MgCl_2_; 25 MgSO_4_; 0.5 KH_2_PO_4_; 2.5 NaHCO_3_; 10 Glucose; 10 HEPES buffer at pH 7.8, osmolarity = 997 mO. Other solutions were made by equimolar substitution of this basic formula. For example, calcium chloride was replaced with barium chloride to enhance the inward L-type calcium current. For the calcium free ASW magnesium was substituted for calcium. Patch pipettes were filled with a solution containing (mM): 500 Aspartate or CsCl_2_; 10 NaCl; 4 MgCl_2_; 3 EGTA; 20 HEPES, 2 Na_2_ATP, 0.2 Na_3_GTP, 0.2 Lucifer Yellow CH (lithium salt), pH and osmolarity were adjusted to 7.4 and 870 Osm mol kg^−1^ H_2_O with KOH and sucrose, respectively.

Pharmacological agents used to block the characterized ionic currents were purchased from Sigma-Aldrich (Gillingham Dorset, England). These included tetrodotoxin (TTX), tetraethylammonium chloride (TEA^+^), 4-aminopyridine (4AP), cesium chloride (CsCl_2_), cobalt chloride (CoCl_2_), barium chloride (BaCl_2_), nifedipine was dissolved in absolute ethanol to make 5 mM stock solutions and stored at 5 °C in the dark. Experiments with nifedipine were carried out in dim light to prevent photo-oxidation. FMRFa was bath applied approximately 10 min after whole cell membrane seal and break-in.

### Statistics

The software package Instat was used for statistical analysis. All data are given as mean ± SE. Where statistical comparisons are made before and after bath application of FMRFa, then the two-tailed paired student’s t-test was employed. If not stated otherwise, data were denoted as statistically significant when *P < 0.05.*

## Results

The morphology of a typical centrifugal neuron located in the inner granule cell layer of the optic lobe as revealed by Lucifer yellow staining through the recording microelectrode is shown in Fig. 1A. These efferent cells have a single axon that crosses the major neuropil area in the plexiform zone and then exits into one of the optic nerve bundles. These centrifugal neurons always give off a number of laterally running branches within the plexiform zone (arrowheads, Fig. 1A) without branching outside this zone. Their long axon can be seen leaving the slice (Fig. 1A). These cells can be provisionally identified visually in living slices of the optic lobe based on their size and position and hence can be readily selected for electrophysiological study and their identity is further verified from their responses to optic nerve bundle stimulation which always evoked an antidromic action current and also by dye filling them through the patch pipette.

**Fig. 1.**
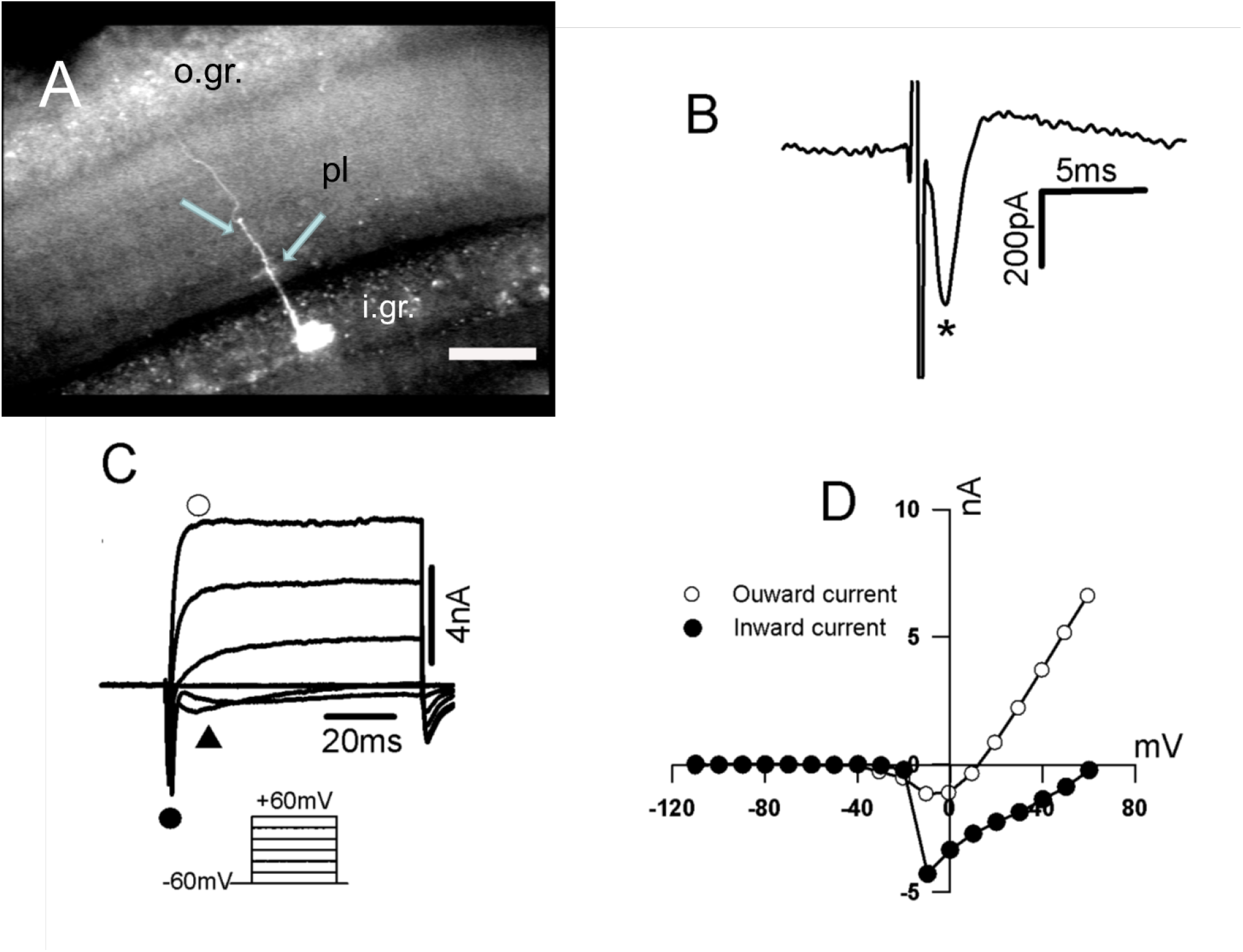
Morphology and overall ionic currents recorded from an identified centrifugal neuron in the optic lobe of cuttlefish. **A)** Lucifer yellow-filled centrifugal neuron with the characteristic numerous fine branches in the plexiform zone (*arrow heads*). o.gr.: outer granule cell layer, pl: plexiform zone and i.gr.: inner granule cell layer. **B)** An evoked antidromic action current resulting from stimulation of the appropriate optic nerve bundle. **C)** Whole-cell currents recorded from a centrifugal neuron. Overall response to a series of voltage steps (with 20 mV increments) from a holding potential of −60 mV is composed of an outward current (*open circle*), a transient inward current (*filled circle*) and a smaller inward current (*filled triangle*). **D)** *I-V* plots of the outward current (*open circles*) and an inward current (filled circles). Horizontal bar: 50 μm (**A**).

In cell-attached mode which provides a way not only to record the activity, but also to stimulate neurons in brain slice preparations. Furthermore, cell-attached recording of action potential currents is an easy type of recording to do because no breaking of the patch is involved, and the seal can be loose (< 1 GΩ; Kondo and Marty, 1998). Using this mode of recording action potential current can be evoked by stimulating the appropriate optic nerve (Fig1. B).

Whole-cell patch clamp recordings were obtained from identified centrifugal neurons, depolarizing pulses of 80 ms duration, from a holding potential of −60 mV, were used to set the membrane potential at voltages ranging from −60 mV to +60 mV with a 10 mV increments which elicited two main categories of currents; a large outward current (*open circle*) and an inward current which had an initial large transient component (*filled circle*) followed by a smaller inward current (*filled triangle*. Fig1. C). The current-voltage (I–V) curves showing the outward current (*open circles*) and the fast inward current *(filled circles,* Fig. 1D).

### Whole-cell outward currents

The large outward current observed in these centrifugal neurons at depolarized potentials was studied using aspartate internal solution in the pipette as well as 1 μM TTX and 2 mM CoCl_2_ in the external solution to block both the sodium and calcium currents respectively. In this study, we found that in 84% of the centrifugal neurons (21 out of 25) displayed two components of the outward current; a transient fast-inactivating component and a sustained component (Fig. 2A). In the remaining 16% (4 neurons), only the sustained component was observed. These two components of the outward current could be separated using either their voltage or pharmacological sensitivities (Connor and Stevens, 1971 a, b). If the cell was held at a holding potential of −40 mV instead of −60 mV, the initial transient component (Fig. 2A) disappeared during the voltage step protocol (Fig. 2B). Subtracting the outward currents recorded at −40 mV from those recorded at −60 mV revealed the shape of the transient or A-current (Fig. 2C). The current-voltage relationships at these two holding potentials (−40 and −60 mV) as well as the isolated transient or A-current (I_A_) are illustrated in Fig. 2D. The addition of 4AP (4 mM) to the bathing solution also abolished the large transient component of the outward current, leaving only the sustained outward current (bold trace, Fig. 2E). The current-voltage relationships for these currents are shown in Fig. 2F. The remaining outward current after either voltage (Fig. 2B) or pharmacological (Fig. 2E) separation has a sustained time course and is similar to the delayed rectifier (I_K_) of other nerve cells. This is confirmed by its reduction (75%) in the presence of TEA (20 mM, Fig. 2E). These results are consistent with the presence of two separate outward currents, which from their kinetics, current-voltage curves, and pharmacological sensitivities, can be characterized as those already reported outward currents in many preparations, namely, the delayed rectifier (I_K_) and the transient A-current (I_A_).

**Fig. 2.**
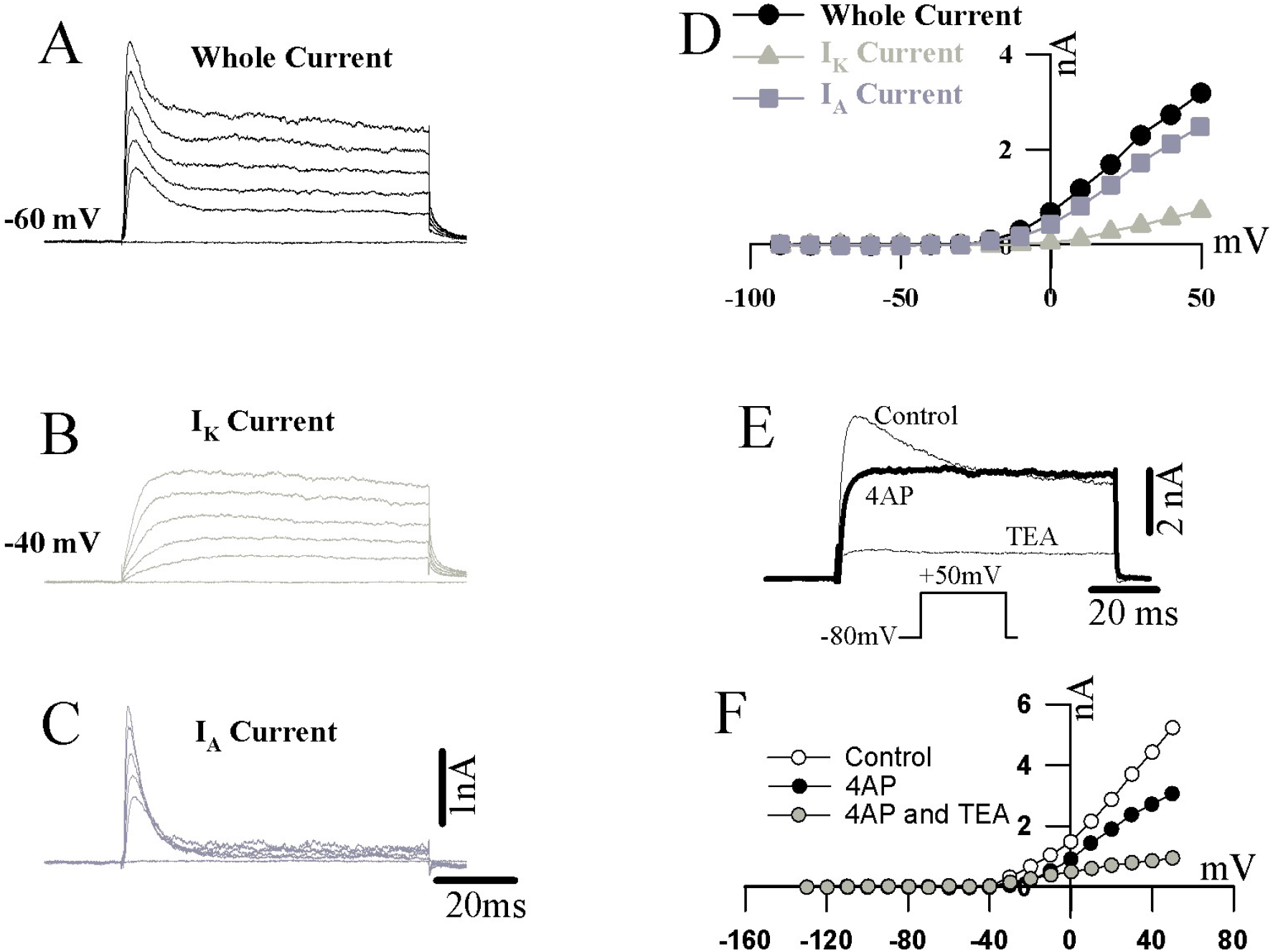
Voltage and pharmacological separation of A-current in an identified centrifugal neuron. **A)** Whole cell outward current in response to membrane depolarization to voltage steps from a holding potential of −60 mV. Total outward current is composed of a transient outward current and a sustained outward current. **B)** Whole cell outward current in response to membrane depolarization to the same voltage steps as in **A** but this time from a holding potential of −40 mV. The A-current is largely inactivated at this holding potential, leaving only the sustained outward current or delayed rectifier (I_K_). **C)** Computer subtraction of **B** and **A** to show the isolated transient outward current or A-current (I_A_). **D** I-V plots of the instantaneous currents 5 ms after the start of the voltage steps for the total outward current (*circles*), the mainly I_K_ current (*squares*) and the isolated I_A_ (*triangles*). **E)** Whole cell outward current in response to membrane depolarization to a voltage step of +50 mV from a holding potential of −80 mV. Total outward current is composed of a transient outward current and a sustained outward current (control). Bath application of 4 mM 4-AP suppressed the transient outward current, leaving only the sustained outward current (4-AP). Bath application of TEA suppressed 75% of this sustained outward current. **F)** I-V plots of the instantaneous currents 5 ms after the start of the voltage steps for the total outward current (*open circles*), the mainly I_K_ current (*filled circles*) and after the application of TEA (*filled triangles*).

### Whole-cell inward currents

Inward currents were studied using cesium chloride for the pipette filling solution to block most of the outward potassium currents we have described above. Under these conditions, two inward currents were observed in centrifugal neurons of the optic lobe slice (Fig. 3). The first one had a transient time course (Fig. 3A, star) and the second had a more sustained time course (Fig. 3A, S). The transient inward current can be blocked by the addition of TTX (1 μM) to the bathing solution, leaving only a sustained inward current (Fig. 3B). TTX is a known blocker of sodium currents in many different animal cell types (Hill, 1992). Subtracting current traces after the application of TTX (Fig. 3B) from those recorded before revealed the TTX-sensitive current (Fig. 3C). The current-voltage relationships (not shown) of this transient inward current appeared at potentials more positive than −50 mV, peaked at around −20 mV, and then starts decreasing. Three lines of evidence indicate that this fast transient inward current is sodium current (I_Na_). Firstly, it was blocked by TTX (Fig. 3B); secondly, it was also blocked in the presence of a sodium-free saline such replacing sodium with lithium or choline chloride (data not shown); and thirdly, this inward current could be progressively inactivated by setting the holding potentials at values more positive than −40 mV. Taken together these data suggest that this current is probably the sodium current responsible for the rising phase of the action potential generation.

**Fig. 3.**
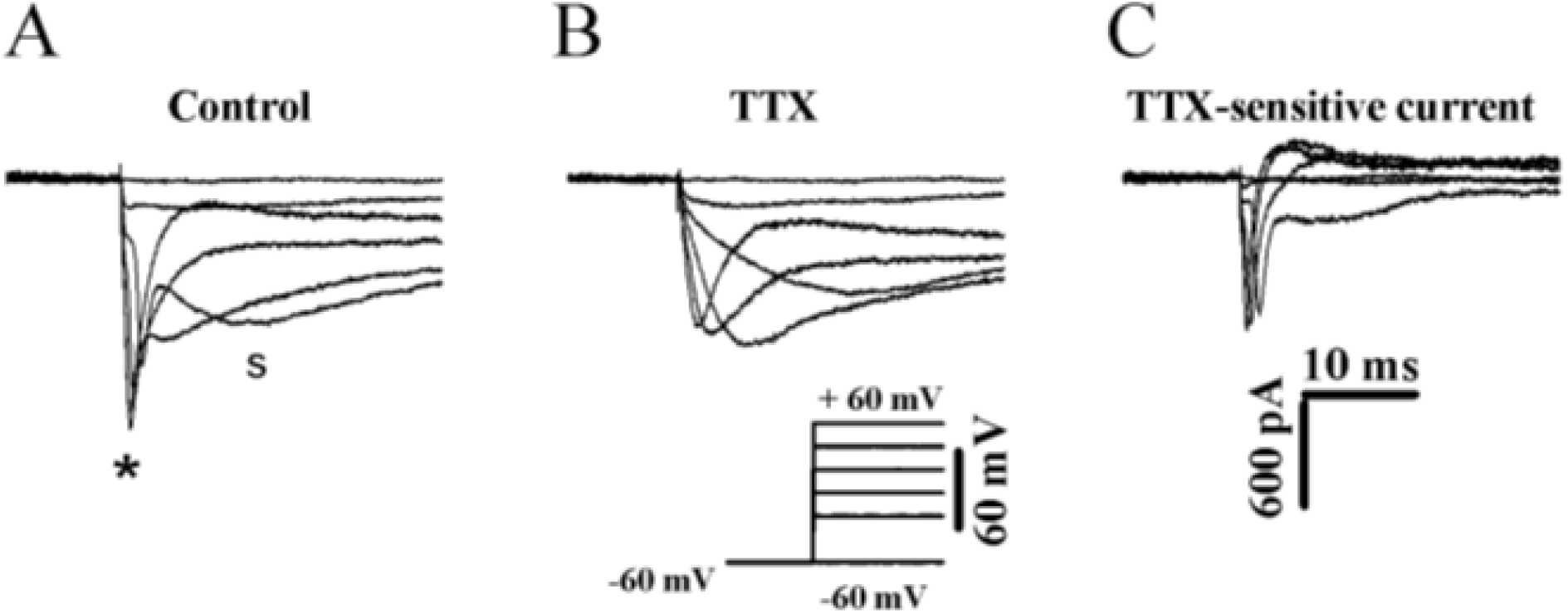
Inward currents recorded from an identified centrifugal neuron. **A)** In the control, the whole-cell currents from this cell obtained in response to a series of voltage steps (*bottom*), from a holding potential of −60 mV. **B)** 5 minutes after bath application of an ASW containing TTX (1μM), the Na^+^ current disappeared leaving only a sustained inward current. **C)** TTX-sensitive current in isolation, which is obtained by computer subtraction of **B** from **A**.

The second inward current had a sustained time course and was not affected either by TTX (Fig. 3B) or by a sodium-free saline (not shown) and could be increased by the substituting barium for calcium in the ASW (Fig. 4A). This current could be totally blocked by the addition of cobalt chloride (2 to 4 mM) to the external solution (Fig. 4B), or cadmium chloride (data not shown) suggesting that it could be carried by calcium ions. Furthermore, this inward current was more importantly blocked by the dihydropyridine antagonist, nifedipine, at a concentration of 5 μM (Fig. 4C) which strongly suggest that this current is probably an L-type calcium current (I_Ca,L_). The current-voltage relationships of this inward calcium current demonstrated that it was rapidly activated by voltage steps more positive than −40 mV and was maintained throughout the voltage step with no sign of inactivation. This sustained inward current achieved a maximum for test steps around 0 or +10 mV and then decreased. In some of our experiments, a third inward current which was also carried by calcium, but can be seen only with test pulses from negative holding potential (V_h_ = −80 mV) had a transient time course compared to the L-type calcium current (Fig. 4D, trace a). The current-voltage relationships plot shown in Fig 4E (filled triangles) show that this current starts to activate at around −50 mV, its amplitude increased progressively with higher depolarizations, reaching a plateau around −30 mV and then began to decrease. This current is usually referred to as T-type calcium current (I_Ca,T_) because of its transient time course. This current is blocked when the holding potential is set at −60 mV or above (Fig. 4D, trace b). This current is similar to the one that have been reported in heart muscles of squid (Ödblom et al. 2000). In these centrifugal neurons, the L-type calcium was encountered invariably. By contrast, the T-type calcium currents were rare and we have only seen them in 5 centrifugal neurons. This transient inward current had similar pharmacological characteristics to that of reported in heart muscle cells of squid (Ödblom et al., 2000; Chrachri et al., 2000), in that it was only partially blocked by 2-5 μM nifedipine. This dihydropyridine antagonist has been shown to block preferentially the L-type calcium current in many preparations (Fox et al., 1987; Wang et al., 1996; Chrachri and Williamson, 1997). However, as for the isolated heart muscle cells, nifedipine also blocks the transient calcium current by about 48.4% (n = 3).

**Fig. 4.**
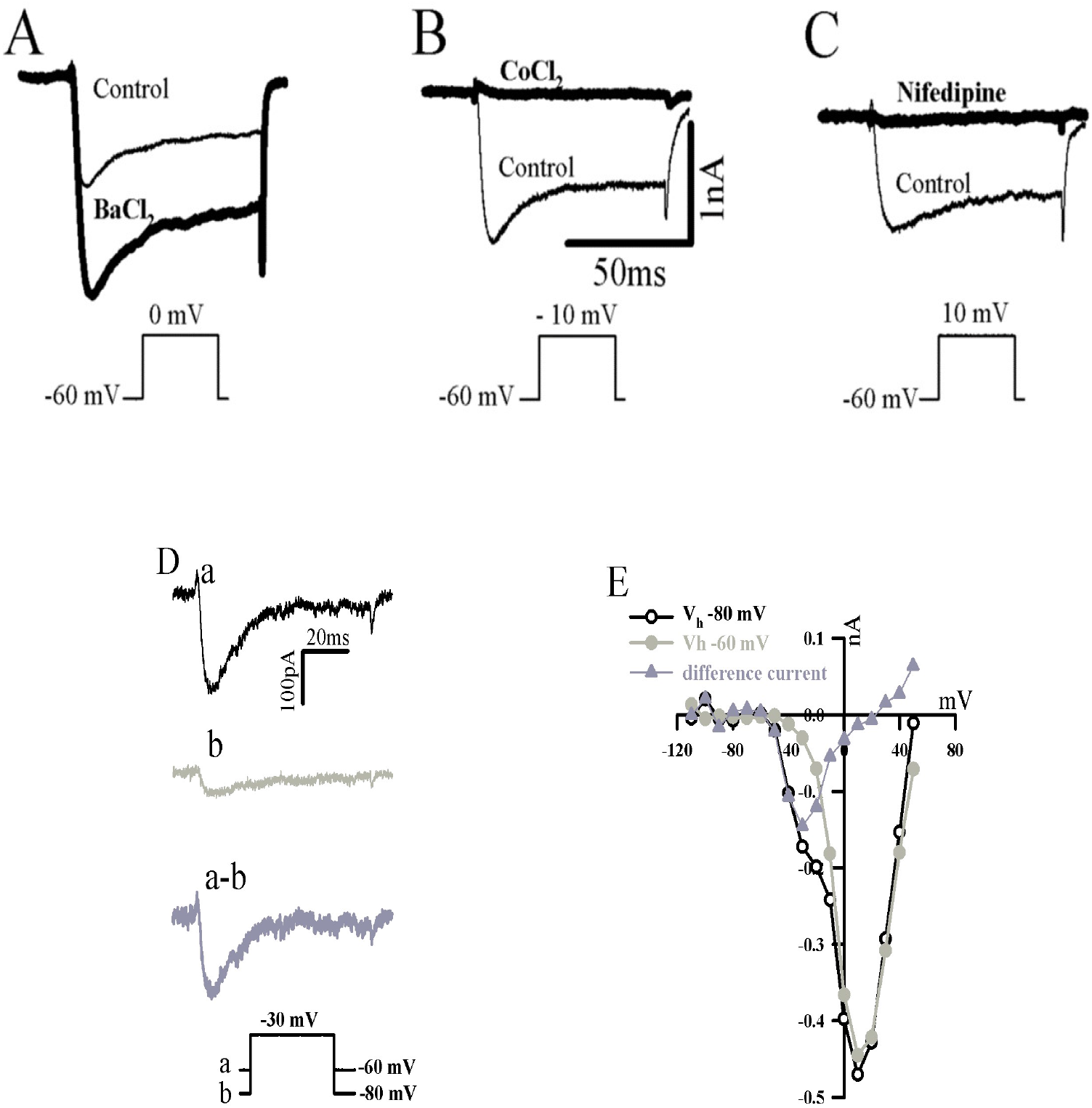
Identification of the L-type calcium current. **A)** Effect of barium chloride on the sustained inward current, whole cell inward current in a centrifugal neuron in response to a voltage step to 0 mV from a holding potential of −60 mV before (*control*) and after (*BaCl2*) substitution of barium for calcium in the external solution which resulted in an increase in the amplitude of the calcium current. **B)** Effect of cobalt chloride at a concentration of 4 mmol l^−1^ on the sustained inward current, whole cell inward current in a centrifugal neuron in response to a voltage step to −10 mV from a holding potential of −60 mV before (*control*) and after (*CoCl_2_*) was added into the external solution resulted in total blockade of the calcium current. **C)** Similarly, nifedipine (5 μmol l^−1^) also suppressed completely this sustained calcium current. **D)** Representative example of membrane currents during a test-pulse to −30 mV from a holding voltage of −80 mV (a) or −60 mV (b); the difference current (a-b) is the T-type 2+ Ca^2+^ current. **E)** Peak current-voltage (I/V) relation of the same centrifugal neuron. For this I/V plot current traces are displayed, when the holding potential was −80 mV (*open circles*), when the holding potential was −60 mV (*filled circles*), and finally the difference current (*triangles*).

### Effect of FMRFa on the outward current

To investigate the effect of the neuropeptide FMRFa on the outward potassium currents in the centrifugal neurons, we bath applied FMRFa (1 μM) to voltage clamped centrifugal neurons and found that this neuropeptide induced a significant reduction of the overall potassium current in 11 out of 15 centrifugal neurons. At a holding potential of −60 mV, the reduction of the potassium currents was significant *(p < 0.0001)* and was about 39.83 ± 7.92 % (n = 11). When the holding potential was set to −40 mV and therefore to study the effect of FMRFa on the delayed rectifier (I_k_) on its own, the reduction of I_k_ by FMRFa was also reduced significantly *(p = 0.0028)* by about 31.27 ± 10.45 % (n = 5). A typical experiment demonstrating the FMRFa-induced reduction of the peak current of both the fast-inactivating and sustained potassium currents is illustrated in Fig. 5. The effect of FMRFa on these two components of the potassium currents was reversible (Fig. 5C). Subtracting the current traces in the presence of FMRFa (Fig. 5B) from the control current traces (Fig. 5A) revealed the FMRFa-sensitive potassium currents (Fig. 5D). In the remaining 4 centrifugal neurons, FMRFa did not appear to have any apparent effect on both components of the outward potassium current (data not shown).

**Fig. 5.**
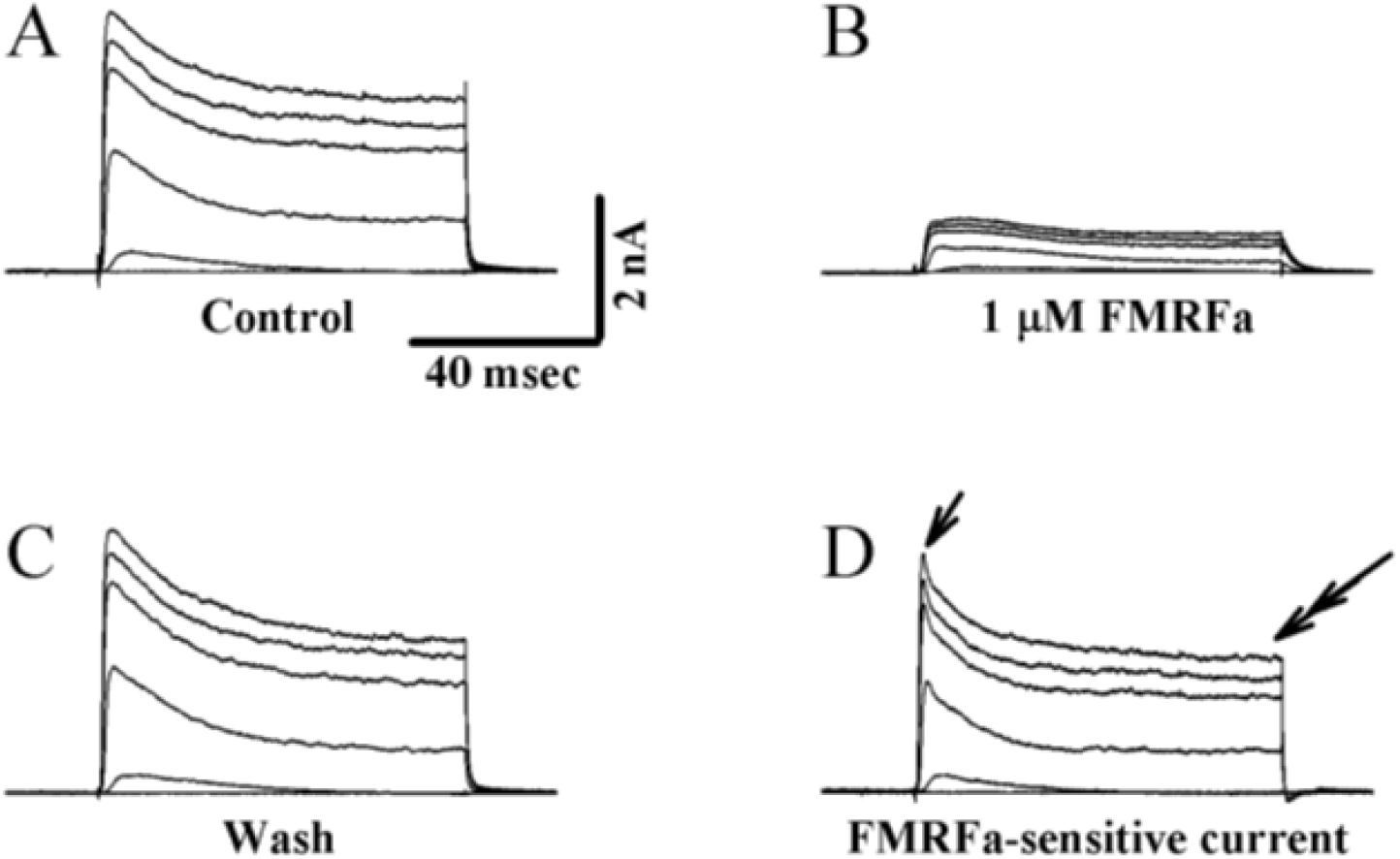
FMRFamide inhibits both the I_A_ and I_K_ in centrifugal neuron. **A)** Current traces recorded in response to membrane depolarization to a voltage step of −60, −20, +20, +50, +60 and +70 mV from a holding potential of −60 mV prior to the application of FMRFa. **B)** Reduction of K^+^ currents by FMRFa (1 μM). **C** Recovery of K^+^ currents after FMRFa had been washed out. **D)** Difference current obtained by subtracting current profiles obtained in the presence of FMRFa **(B)** from those obtained before the application of FMRFa **(A)** demonstrates that FMRFa blocked both the fast transient component, I_A_ (*arrow head*), well as the sustained and slowly inactivating potassium current, I_K_ (*double darrow head*).

### FMRFa-mediated attenuation of the inward sodium current (I_Na_)

Bath application of FMRFa (1 μM) seems to induce a reduction in the magnitude of this inward sodium current. Fig. 6A illustrates a typical example, 5 minutes after bath application of FMRFa the amplitude of the sodium current was reduced by about 28.8% (Fig. 6A, grey line). 8 minutes after the addition of FMRFa, the amplitude of I_Na_ was reduced even further, this time by about 44.44% (Fig. 6A, thick line). The effect of FMRFa on the I_Na_ is also illustrated in Fig. 6B, where the current-voltage relationship demonstrates that FMRFa reduces I_Na_ over most of the voltage ranges. Pooled data from 3 experiments demonstrated that FMRFa induced a significant (*p = 0.01*) decrease of the sodium current by about 41.74 ± 5.94%.

**Fig. 6.**
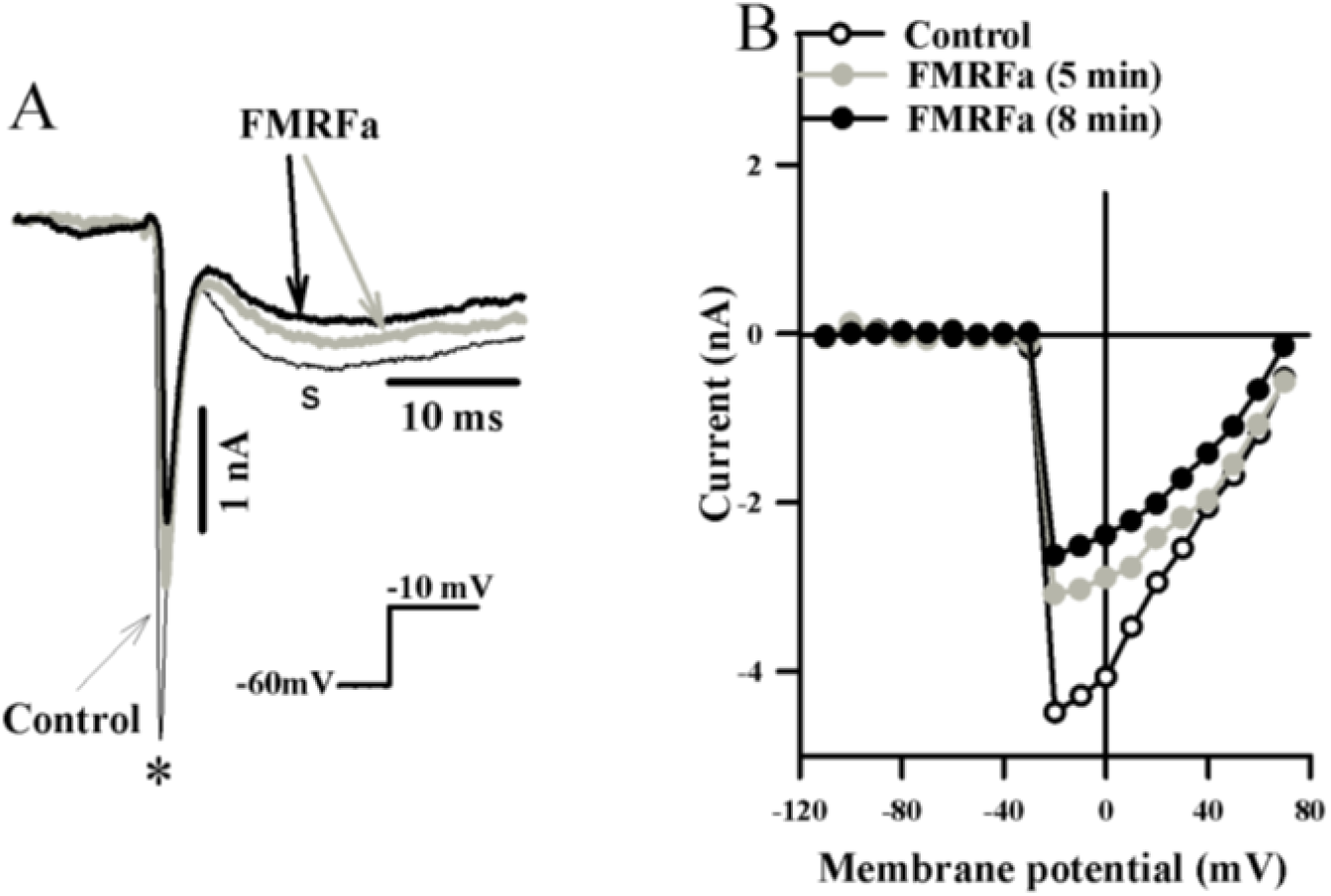
FMRFamide-mediated inhibition of I_Na_ and I_Ca,L_ in a centrifugal neuron. **A**) Current traces recorded in response to membrane depolarization to a voltage step of −10 mV from a holding potential of −60 mV before (*control*) and after (*FMRFa*) bath application of 1 μM FMRFa, first after 5 minutes (*grey trace*) and then 8 minutes (*dark trace*). These currents trace not only show that FMRFa inhibited I_Na_ (*star*), but also I_CaL_ (*triangle*). **B)** I-V relationship of I_Na_ before (*open circles*), 5 minutes and then 8 minutes after (*grey* and *filled circles,* respectively).

### Effect of FMRFa on L-type and T-type calcium currents

Fig. 6, not only demonstrates that FMRFa reduced the peak amplitude of the I_Na_ (star), but also decreased that of the sustained L-type calcium current (triangle). Unlike in the case of potassium currents, bath application of FMRFa (1 μM) always resulted in reducing the calcium current. The magnitude of the peak I_Ca,L_ was reduced significantly (*p = 0.0002*) by about 64.92 ± 5% (n = 8). An example is illustrated in Fig. 7A (right panel), where the inhibition of I_Ca,L_ was about 52%. The current-voltage curves show that FMRFa reduction of the amplitude of I_Ca,L_ is over most of the voltage ranges. However, the reduction induced by FMRFa decreased for steps to depolarized voltages more positive than +10 mV (Fig. 7B). This figure also demonstrates that FMRFa (1 μM) didn’t have any noticeable effect on the T-type calcium current (Fig. 7A, left panel). The current/voltage plot provides further evidence that FMRFa selectively affects the amplitude of I_CaL_, but not the I_CaT_ (grey circles, Fig. 7B). Figure 8 illustrates the time course of the onset of the inhibition of I_CaL_ by FMRFamide and the subsequent recovery by washing, and demonstrates that the inhibition was relatively rapid and fully reversible.

**Fig. 7.**
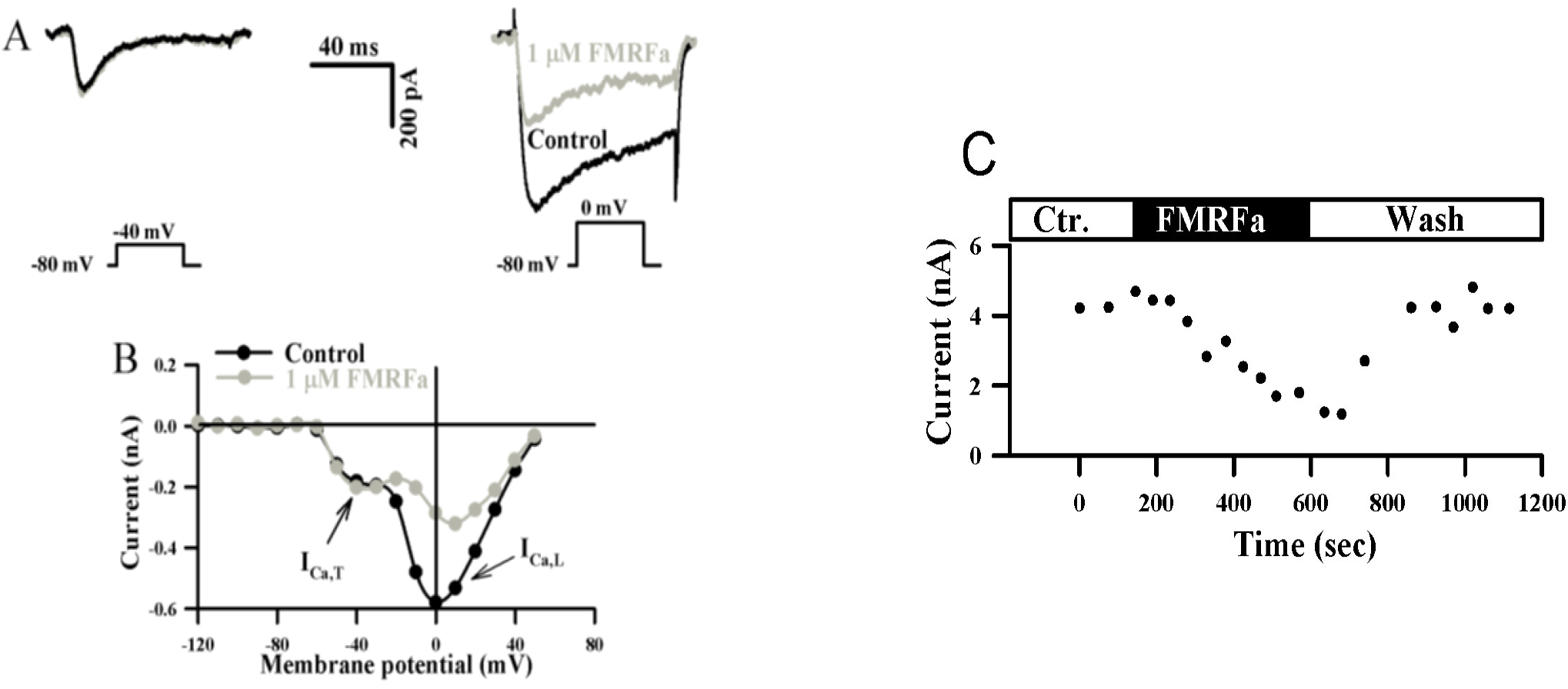
Effect of FMRFamide on calcium currents in centrifugal neuron. **A)** Left traces are current traces recorded in response to membrane depolarization to a voltage step of −40 mV from a holding potential of −80 mV showing that FMRFa had no apparent effect on the transient component of the calcium current. Right traces are current recorded in response to membrane depolarization of the same centrifugal neuron to a voltage step of 0 mV from a holding potential of −80 mV before (*black trace*) and after (*grey trace*) the application of FMRFa demonstrating that this neuropeptide decrease the amplitude of the I_CaL_. **B)** I-V relationship of both calcium current under control conditions (*black circles*), and after the application of FMRFa (*grey circles*). **C)** Time course of the onset, and recovery from, the effect of FMRFamide on I_CaL_.

## Discussion

This study has characterized the electrical properties in identified centrifugal neurons in the optic lobe slice preparation of the cuttlefish, *Sepia officinalis,* using whole-cell patch clamp we documented the presence of five voltage-sensitive ionic currents and studied their modulation by the neuropeptide FMRFa. These voltage-activated currents comprised; tree inward currents; one carried by sodium (I_Na_) and the two others by calcium (I_CaL_ and I_CaT_), and two outward currents that were selective for potassium (I_A_, I_K_). The effect of the neuropeptide, FMRFa, on these currents is also discussed.

### Na^+^ current

The rapid inward current is activated immediately after the onset of the command pulse, reaches a peak in few milliseconds, and inactivated within 10 ms. As in many other excitable tissues (Lo and Shrager, 1981; Neher, 1971; Lasater, 1986), this fast inward current is carried by sodium, in that it is blocked when the optic lobe slice is perfused with a sodium-free ASW, it is also totally blocked by TTX (1μM) and is inactivated when the cells is held at values more positive than −40 mV. This I_Na_ was present in all of the centrifugal neurons we have recorded from (without exception). The presence of such sodium current in these centrifugal neurons was not surprising because these neurons send their axons a long distance towards the retina (Saidel, 1979) and therefore it is possible that these centrifugal neurons will need action potentials to carry their information towards the photoreceptors located in the retina. Whole-cell patch clamp recordings from other cell types within the optic lobe (i.e. medulla and amacrine) demonstrated that these neurons do not extend their axons towards the retina, and that only a few of them displayed sodium currents it is present in only about a third of them (Chrachri and Williamson, unpublished data).

### Ca^2+^ currents

A vast amount of investigations have been carried out on the voltage-dependent calcium currents in a variety of neuron and muscle cells (Liu and Lasater, 1994; Ödblom et al., 2000). In the present investigation we have identified two more inward currents, one of which have a sustained time course and the other have a transient time course. Both of them were carried by calcium ions (Fig. 4). On the basis of its kinetics, ion specificity and pharmacological sensitivity, the sustained inward current resembles to the high voltage-activated calcium (HVA), or L-type calcium current reported in a variety of other cell types (Carbone and Lux 1984,; Fox et al., 1987). The other voltage-dependent calcium current is similar to the transient, low-threshold (LVA) or T-type (Nowycky et al., 1985) which only appeared when neurons were held at potentials more negative than −60 mV. This current had similar kinetics and pharmacological characteristics to that reported in squid isolated heart muscle cells (Ödblom et al., 2000; Chrachri et al., 2000), in that it showed higher selectivity to blockade by nickel chloride than the L-type calcium current (data not shown) which is similar to reports in other preparations (Mitra and Morad, 1986; Hagiwara et al., 1988; Wu and Lipsius, 1990). It was also partially blocked by nifedipine which has been shown to have a more specific effect on the L-type calcium current (Scott et al., 1991), but similar results have been reported in atrial cells (Bean, 1985), retinal ganglion cells (Liu and Lasater, 1994) and olfactory bulb neurons (Wang et al., 1996).

### K^+^ currents

Under voltage-clamp conditions, when centrifugal neurons were depolarized to voltages more positive than −30 mV from a holding potential of −60 mV, two outward potassium currents were detected, the delayed rectifier or I_K_ and the A-current or I_A_. These two time- and voltage-dependent K^+^ currents were distinguishable both by their pharmacological sensitivity to 4AP, and by their voltage-dependent inactivation (Connor and Stevens, 1971 a, b; Thompson, 1977). Similar results have been reported in other cephalopod cells (Llano and Bookmanm, 1986; Lucero et al., 1992; Chrachri and Williamson, 1997). TEA at a concentration of 20 mmol l^−1^ blocked most of the delayed rectifier K^+^ currents in centrifugal neurons. These concentrations of TEA are relatively very low compared to those needed to block the similar I_K_ in sensory hair cells of the squid statocysts (Chrachri and Williamson, 1997), the squid giant axons (Tasaki and Hagiwara, 1957) and isolated heart muscle cells (Ödblom et al., 2000) where very high concentrations of TEA were needed. The delayed rectifier activates at potentials more positive than does the A-type current and show no appreciable inactivation during a depolarizing voltage step also contribute to action potential repolarization (Saito and Wu, 1991). The effects of blockade of delayed rectifier in centrifugal neurons by TEA are consistent with the pharmacology of other delayed-rectifier subtypes of K^+^ channels (see Hille, 1992).

### FMRFa-mediated effect on the centrifugal neurons of the optic lobe

We have demonstrated that the neuropeptide, FMRFa, significantly reduced the inward Na^+^ current in the centrifugal neurons of the optic lobe, the effect was voltage dependent, it lasts as long as the peptide is present and finally the reduction of Na^+^ current by FMRFa is partially reversible. Furthermore, This FMRFa-induced inhibition was also seen as a reduction of the antidromic action current following stimulation of the appropriate optic nerve bundle (not shown). Modulation of Na^+^ current is likely to be important in the regulation of the centrifugal neuron’s excitability. The reduction of the I_Na_ by FMRFa will increase the threshold of the action potential and so contribute to the inhibitory effects of FMRFa on the excitability of the cell, possibly resulting in suppressing on-going firing activity as well as preventing the generation of new discharges. Inhibition of this current by FMRFa has been shown in other molluscan preparations, the peptidergic caudo-dorsal cells of the mollusc *Lymnaea stagnalis* (Brussaard et al., 1991).

Another action of FMRFa is to attenuate the voltage-dependent sustained calcium current. Attenuation of this type of calcium current by FMRFa has also been described in a number of other types of cells (Kramer et al., 1988; Man-Song-Hing et al., 1989; Yakel, 1991; Chrachri et al., 2000). However, this neuropeptide didn’t have any effect on the T-type calcium current. Similar selectivity for the FMRFamide-induced reduction of the calcium current has been reported in neurons of *Aplysia californica* (Brezina et al., 1987), in isolated heart muscle cells of squid (Chrachri et al., 2000). It is probably not surprising that FMRFa affected I_Ca,L_ but not I_Ca,T_ in centrifugal neurons. It has been reported that this neuropeptide reduced the amplitude of spontaneous excitatory postsynaptic currents (sEPSCs) in centrifugal neurons. However, when these sEPSCs occurred in bursting mode, FMRFa did change the amplitude of sEPSCs without any significant modulation of the frequency their rhythmic activity (Chrachri unpublished data). This transient current has been proposed to control rhythmic membrane oscillations central neurons (Llinas and Yarom, 1981) and therefore the lack of effect of FMRFa on I_CaT_ might have been expected.

We have also presented evidence that application of FMRFa evoked a partial blockade of both characterized outward; the I_A_ as well as the I_kv_ currents in these morphologically identifiable centrifugal neurons of the optic lobe.

### Physiological relevance of the action of FMRFa

The reduction of the amplitude of the L-type calcium current in the centrifugal 2+ neurons would be a relevant physiological action for FMRFa. Ca^2+^ channels in neurons are frequent targets of neurotransmitter modulation, with suppression or 2+ enhancement of Ca^2+^ channel activity being a common outcome, albeit *via* different mechanisms (Gerschenfeld et al., 1989; Yakel, 1991). The fact that this neuropeptide affect the ionic currents (this study) and the postsynaptic currents of these efferent neurons (Chrachri and Williamson, 2003) would undoubtedly suggest a role for the retinal cells which are under the control of these centrifugal neurons (Saidel, 1979). There is a wide distribution of FMRFa like immunoreactivity in the optic lobe of cephalopods (Suzuki et al., 2002; Chrachri and Williamson, 2003). Thus the FMRFa-induced effects on the centrifugal neurons we have reported in this paper may probably mirror the endogenous modulation of the centrifugal neuron by FMRFamidergic nerve fibres. Using behavioral studies it has been demonstrated that the optic lobe of *sepia officinalis* may provides a system for coding, sorting and decoding the visual input to produce a relevant behaviour (Chichery and Chanelet, 1976). The fact that the application of FMRFa induced not only a modulation of ionic currents (this report), but also appears to act presynaptically to modulate both spontaneous excitatory and inhibitory currents in the same preparation (Chrachri and Williamson, 2003) could suggest that this neuropeptide may play a crucial role in visual processing. Similar roles for FMRFa in visual processing have been described in the locust optic lobe (Rémy et al., 1988) and in fish (Wang et al., 2000).

## List of symbols and abbreviations

o.gr.: outer granule cell layer
i.gr.: inner granule cell layer
pl: plexiform zone
4AP: 4-aminopryridine
TEA: tetra-ethyl ammonium
TTX: tetrodotoxin

